# Robust and Efficient Coding with Grid Cells

**DOI:** 10.1101/107060

**Authors:** Lajos Vágó, Balázs B Ujfalussy

## Abstract

The neuronal code arising from the coordinated population activity of grid cells in the rodent entorhinal cortex can uniquely represent space across large distances but the precise conditions for efficient coding are unknown. Here we present a number-theoretic analysis of grid coding and derive an upper bound on the distance that a population of grid cells can represent without error. We show that in the absence of neuronal noise, the capacity of the system would be extremely sensitive to the choice of the grid periods. However, when the accuracy of the representation is limited by neuronal noise, the capacity becomes gradually more robust against the choice of grid scales as the number of modules increases and remains near optimal even for random scale choices. Our study reveals that robust and efficient coding can be achieved without parameter tuning in the case of grid cell representation.

## 1. Introduction

Optimising neuronal systems for efficient processing and representation of information is a key principle for both understanding and designing neuronal circuits (Sterling & Laughlin, 2015), but recognising that a particular neuronal phenomena reflects an optimisation process is often difficult. Grid cells in the medial entorhinal cortex have been suggested to represent spatial location by their spatially periodic firing fields near optimally (Burak et al., 2006; Fiete et al., 2008; Mathis et al., 2012a,b). However, it remained controversial whether the robustness and efficiency of the grid cell code is the result of the precise tuning of the grid parameters (Wei et al., 2015; Stemmler et al., 2015; Mosheiff et al., 2016) or the performance of the system is relatively insensitive to the actual parameter settings (Mathis et al., 2012a,b; Towse et al., 2014).

Grid cells are spatially tuned neurons with multiple firing fields organised along the vertices of a triangular grid (Figure 1a; Hafting et al. 2005; Moser et al. 2014). Grid cells of any particular animal are organised into functional modules (Barry et al., 2007; Stensola et al., 2012): cells within a module share the same grid scale and orientation, but differ in the location of their firing fields, i.e., their preferred firing phase within the grid period (Figure 1a). Modules form the functional units of the grid representation: The joint activity of all (possibly hundreds of) cells within each module is captured by the (two dimensional) *phase* of the given module (Figure 1b; Sreenivasan & Fiete 2011; Yoon et al. 2013) and the relationship between different cells from the same module remains stable across different environments (Fyhn et al., 2007) or after environmental distortions (Stensola et al., 2012). A given spatial location is represented by the phases of the different modules (*phase vector*). The representations are unique up to a critical distance above which the coding becomes ambiguous: the phase vectors, and hence the firing rates of all grid cells, become (nearly) identical at two separate physical locations (Figure 1c).

**FIGURE 1.**
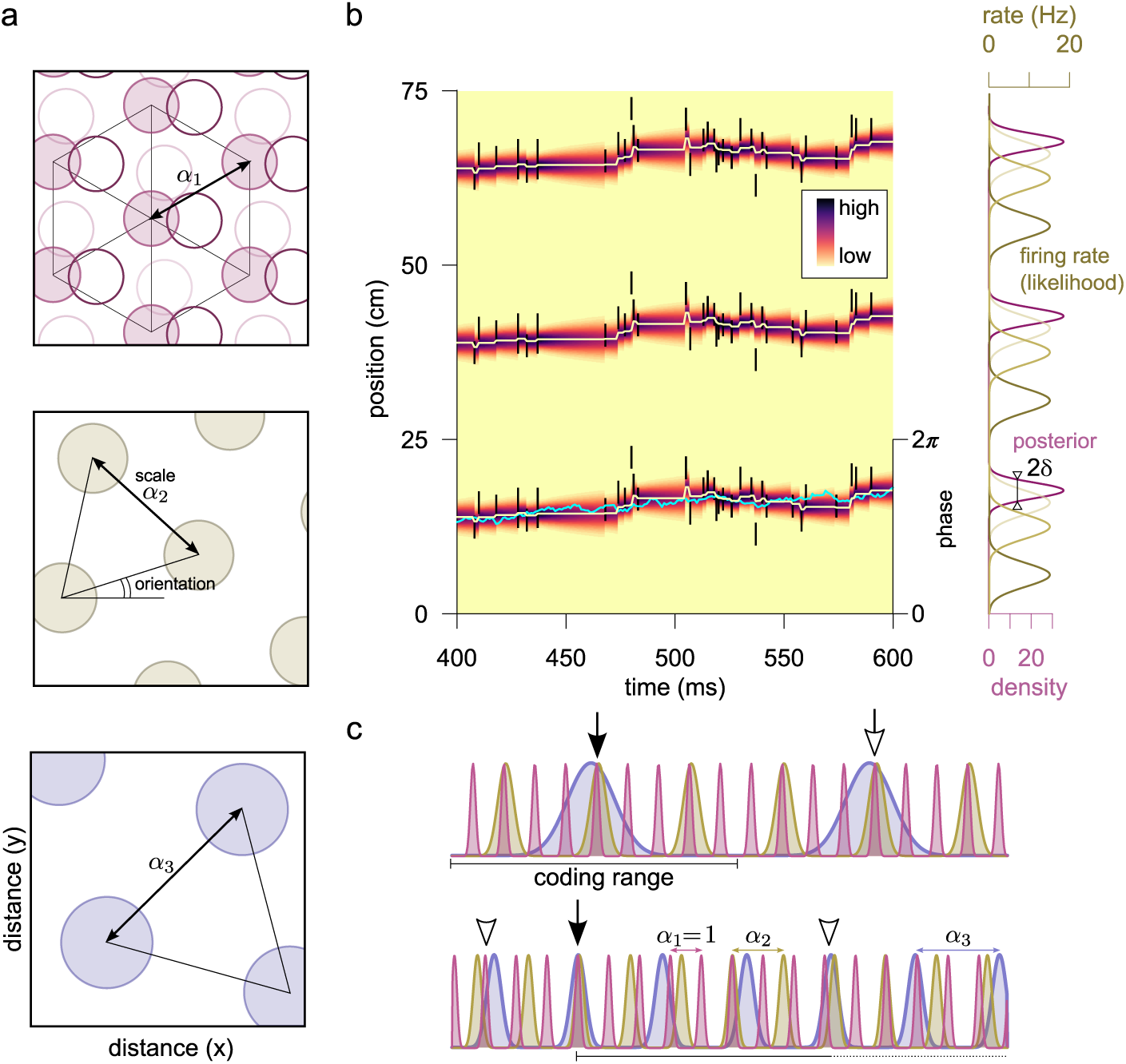
Coding with grid cells. (a) Schematic firing fields (circles) of two-dimensional grid cells as function of spatial position. Grid cells are organised into modules: Cells from the same module share the orientation and scale parameter but differ in their spatial phase (top, shades of purple). Different modules have different scale and orientation (top to bottom). (b) Spiking of grid cells (black ticks, each spike is shown three times, at the maxima of the cells’ firing rate) from a single module represents the movement of the animal (light-blue line) in a 1 dimensional environment. Since the firing rate of the cells (right, olive) is periodic, the posterior distribution (left: colormap, right: purple) is also periodic. The accuracy of the representation, quantified by the width of the posterior distribution (*δ*, right) changes over time around a typical value that depends on the scale and the firing rate of the module. (c) Grid cell coding schemes. The location of the animal (filled arrow) is jointly encoded by the posterior of the different modules in both nested (top) and modulo arithmetic (bottom) codes. Empty arrows indicate locations with large interference between the modules.

Depending on the magnitude of the critical distance compared to the largest grid scale, two complementary coding schemes have been proposed for grid cells (Figure 1c): In *nested coding* (Mathis et al., 2012a; Wei et al., 2015; Mosheiff et al., 2016) the coding range is set by the module with the largest scale and the smaller modules refine the position coding. In a *modulo arithmetic* (MA) code (Fiete et al., 2008; Sreenivasan & Fiete, 2011) the coding range can be substantially larger than the scale of the largest module and the grid scales have similar magnitudes.

It has been demonstrated that grid cells organised in a nested coding scheme can unambiguously represent spatial locations up to a distance that is exponentially high in the number of modules (Mathis et al., 2012b; Wei et al., 2015; Mosheiff et al., 2016). However, an efficient nested code demands accurate tuning of the scale parameters as well as the number of neurons in a given module, and even then its coding range is constrained by the largest grid scale. It remained unclear whether the sensitivity to its parameters is a general property of all exponentially strong grid cell codes or it is specific to the nested coding scheme. In particular, it is not known under what conditions the MA coding system can achieve exponential capacity, and how robust is the capacity to the choice of the grid periods or neuronal noise.

Here we develop a novel approach to study the capacity of the grid coding system that is based on Diophantine approximations, i.e., approximation of real numbers by rational numbers. First, we apply the technique to study coding with two grid modules. We show that the capacity of the system is extremely sensitive to the number theoretic properties of the scale ratio between the modules. Next, we generalise our approach to the case of multiple modules, and show both analytically and numerically that the exponential capacity of the grid cell coding system can be achieved under very general conditions. Finally, we demonstrate that when the coding range is constrained by neuronal noise, the capacity of the system is extremely robust for the choices of the scaling of the modules.

## 2. Results

We investigate grid cell population code along a linear trajectory as the one dimensional results extend to two (or higher) dimensions without difficulty (Fiete et al., 2008; Mathis et al., 2012a). As the joint firing pattern of cells from the same module is periodic (with the period being equal to the scale of the grid, Figure 1b), it is convenient to prescribe that, without loss of generality, at the spatial origin all modules are in their 0 phase. Then the phase of module *i* is determined by the spatial position *x* of the animal as *ψ*_*i*_(*x*) := (*x* mod *α*_*i*_)/*α*_*i*_ where *α*_*i*_ is the scale of the module with *α*_0_=1, which means that distances are expressed in the unit of the smallest grid period.

Spikes of the neurons in module *i*, s_*i*_, represent the spatial location of the animal with a relative error *δ*. *α*_*i*_ which can be interpreted as the width of the (periodic) *posterior probability distribution P*(*x*|s_*i*_) (Figure 1b). For an ideal observer this posterior distribution quantifies how much a given spatial location is consistent with the observed spike pattern. Naturally, the width of the posterior depends on several factors, most importantly on the number of neurons observed in a given module and on the scale of the modules relative to the typical speed of the animal (Mosheiff et al., 2016). In the methods 4.1 we estimate that the realistic range of *δ* is 0.01 ≤ *δ* ≤ 0.2 which is also consistent with the estimates of Mosheiff et al. (2016).

We assume that the modules are conditionally independent given the location of the animal, and hence position decoding, or representation, can be implemented by an ideal observer independently reading out the spikes, s*_i_*, emitted by the different modules: *P*(*x*|s) = Π*_i_ P*(*x*|s*_i_*). Using numerical simulations (methods 4.1) we found that the module-posteriors can be accurately described as being periodic Gaussian functions (Figure 1b, right). In the following we will treat them as idealised periodic bumps with a width parameter *δ* When loosely talking about interference between the grid modules we refer to the interference between these periodic posterior distributions *P*(*x*|s*_i_*).

### 2.1. Interference between the modules

We illustrate the effect of the scale ratio on the joint firing pattern of the grid cells, and hence on the performance of the grid system for two modules in Figure 2. The goal of avoiding interference between successive modules motivated the choice of irrational scale ratios (Figure 2B, Sreenivasan & Fiete 2011). In the noiseless case, when *δ* → 0, any irrational scale ratio guarantees an infinite capacity for the grid cell coding system even with two modules. However, neurons are noisy, and therefore interference may occur even with irrational scale ratios. It is unclear how irrational numbers compare in terms of robustness against interference.

**FIGURE 2.**
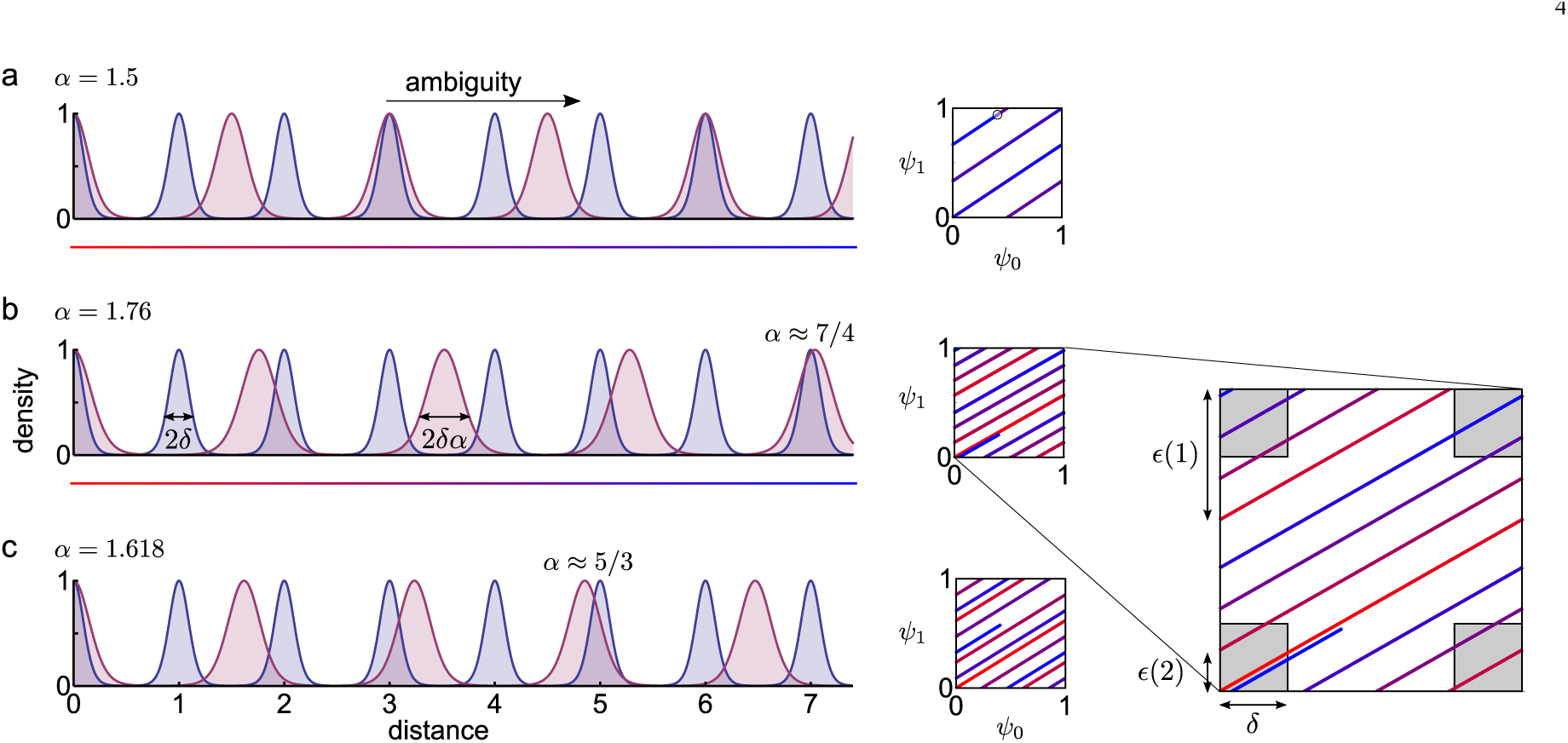
Interference depends on the choice of the scales. (a) Interference with rational scale ratio. Left: Representative posteriors (*P*(*x*|s)) for two modules with scale 1 and *α* = 3/2. Encoding becomes ambiguous at distance 3 from the origin where perfect interference occurs (3 = 2*α*). Right: Phase plot of the two modules, with the colour (red to blue) encoding the distance from the origin (see the coloured line below the left panel). Perfect interference occurs when the phase-curve overlaps with itself. (b) Interference with *α* = 1.76…, which is close to 7/4 and therefore leads to strong interference at distance 7. Right: Interference occurs when the distance between two neighbouring segments of the phase curve becomes smaller than the limit set by the neuronal noise, Graphically, interference corresponds to the phase curve intersecting one of the squares of side *δ* around the origo (inset, grey) or the phase difference, *∊*(*ℓ*) being smaller than 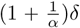 (c) Interference with *α* = *σ* ≈ 1.618, which is the golden ratio. Interference still becomes stronger at larger distances, (e.g. at distance 5, since 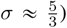 Interference of grid codes is relatthen it would also confuse the origin withed to the approximation of irrationals with rational numbers having small denominators (see text for further details). Right: Interference is inevitable since the phase space has a limited volume.

To quantify interference between two modules with period 1 and *α* up to a maximum distance *L* from the origin, it is enough to check interference in all positive integer distances *ℓ* ≤ *L*, where the phase of the first module, *ψ*_1_(*ℓ*), is 0. Low interference between the modules requires their phase difference *∊*(*ℓ*) at integer distance *ℓ* being large, or equivalently, the phase of the second module being different from 0:

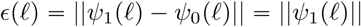

where ∥ can not be the phase difference in general, it is the phase difference at integer distance *ψ*∥ means distance from the nearest integer. From now on we will loosely call *∊*(*ℓ*) the *phase difference*^3^.

If interference is low for all positive integers *ℓ* ⩽ *L*, then not only the coding of the origin will be unambiguous, but all positions in the interval [0, *L*] will be distinguishable by the grid code. Indeed, if the grid code was ambiguous confusing spatial locations *x*_1_ and *x*_2_, then it would also confuse the origin with |*x*_1_ − *x*_2_| as well, since the phase differences of each module are the same between 0 and |*x*_1_ − *x*_2_| and between *x*_1_ and *x*_2_ (Figure 2bc, right)4. Therefore, strong grid codes require uniform coverage of the phase space across arbitrary distances (Sreenivasan & Fiete, 2011).

To analyse the coding properties of the grid cell system, we follow the same three logical steps both in the two module and in the multi-module case (Figure 3). First, we show the existence of an upper bound on how the phase difference *∊*(*ℓ*) between the modules decreases with the distance. Intuitively, this upper bound expresses the fact that interference between the modules necessarily becomes stronger at larger distances. Second, we demonstrate that for appropriately chosen scale ratios a lower bound on the phase difference also exists. For these scale choices catastrophic interference is avoided until a critical distance, that depends on the noise level in the system. Importantly, the slope of the bounds depends only on the number of modules, but not on the choice of the scale parameters. Therefore the efficiency of the scale choices can be characterised by the offset parameter, *c_α_*, of the lower bound. Thus, our third step is to calculate *c_α_* for various choices of the scale parameter *α*.

**FIGURE 3.**
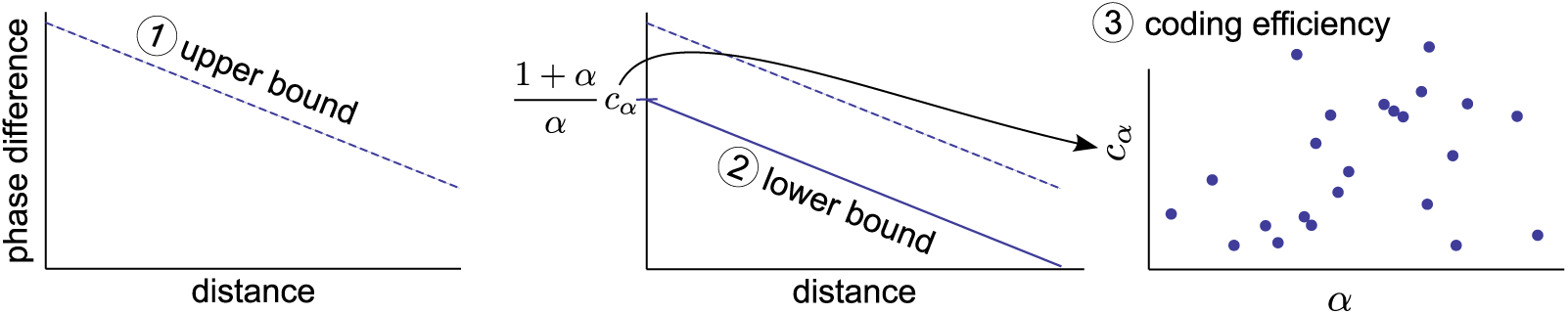
Logical steps of the argument. First, we provide an absolute upper bound on the phase difference in the function of the distance, that is linear in log-log scale (left). Second, we show, that for certain scales a lower bound also exists (middle). Third, we characterise the efficiency of the scales (*α*) by their offset, *c_α_* (right).

To better illustrate our approach, we first consider only two grid modules. Our analytical derivations provide an estimate for the asymptotic performance of the system that is valid in the low-noise limit. The main advantage of our approach is that it provides strict bounds on the achievable coding efficiency that can be compared with the efficiency at realistic noise levels, two or more modules alike.

### 2.2. Coding is extremely sensitive to the scale ratio with two modules

We can formalise the problem of interference between two modules as having a pair of integers *k* and *ℓ* with *ℓ* ≈ *kα* meaning that module 2 (with scale *α*) is close to being in phase 0 at distance *ℓ*, which would cause ambiguity between the coding of the spatial point *ℓ*
and the origin. This is formally identical to the number theoretic question of the approximability of the scale *α* ≈ *ℓ*/*k* with rationals having numerator *ℓ* < *L*, also known as Diophantine approximations (Figure 2bc).

A classical result in number theory (Hurwitz, 1891) states that for all irrational numbers *α* > 1 there are infinitely many relative primes *k*, *ℓ* such that the error of the approximation, defined as

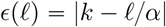

is smaller than the upper bound:

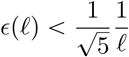

Note that Equation 2.2 expresses the same quantity as Equation 2.1 since*ψ*_2_(*ℓ*) = [*ℓ*/*α*] mod 1, where the approximation error corresponds to the phase difference between the modules (Figure 2bc).

Applying to the grid cells, Hurwitz’s theorem provides an upper bound on how the phase difference between the modules shrinks with the distance. Specifically, the theorem states that the phase difference is guaranteed to shrink at least as *∊*(*ℓ*) ∝ 1/*ℓ* (Figure 4a, dashed line) implying that on the long run interference can not be avoided no matter how carefully we choose *α*. This is a fundamental upper bound on the efficiency of coding with grid cells.

**FIGURE 4.**
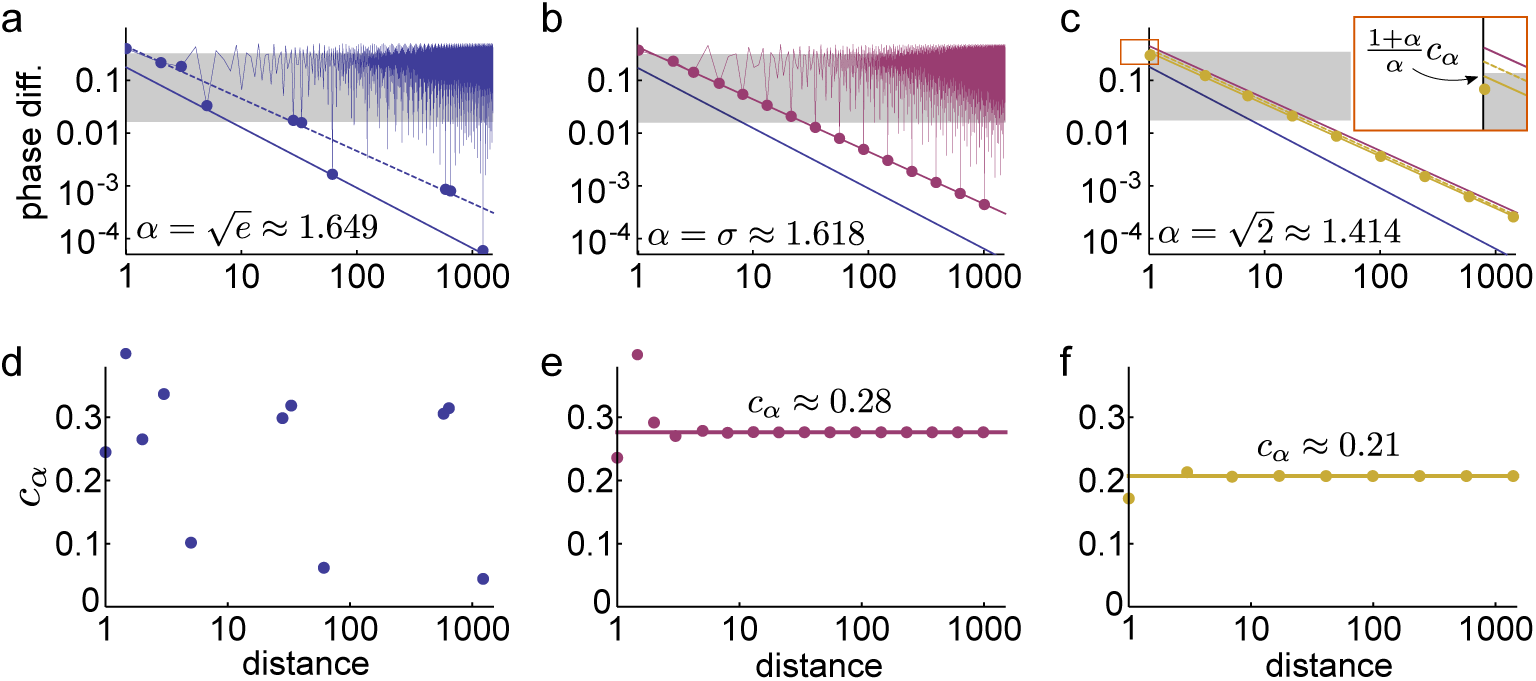
Coding efficiency in two modules with different scale ratios. (a-c) The minimal phase difference always decreases in the function of distance. The zig-zag line indicates the phase difference (PD) at all integer distances, circles indicate record low PD. Dashed line shows the theoretical upper bound of the PD, solid line shows the numerical fit on the lower bound (with slope being −1 and allowing finitely few exceptions at low ℓ). Note, that the lower and the upper bound coincides in b. Also note the 1/*ℓ* scaling of PD for algebraic scale ratios (b-c). Grey shading indicates the range of PD smaller than noise, (1 + *α*)*δ*. **(d)-(f)** The value of *c*_α_(*ℓ*) (Equation 2.6) for different distances and scale ratios. *c_α_* is the highest constant under which there are only finitely few values of *c_α_*(*ℓ*) at small distances. The value of *c*_α_ is slightly higher for the golden ratio (e) than for √2 (f) and much larger than for non-algebric numbers (d, *α* = √e, *c_α_* = 0).

Whether the phase difference is small or large depends on the noisiness of the two modules, *δ* and *αδ*. The representation of the position becomes ambiguous if the phase difference is smaller than the noise in the two modules: 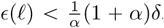 where the 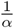 factor comes from the conversion from distance to phase. Therefore, ambiguity does occur at distance *ℓ* ± *δ* from the origin for some *ℓ* if the noise in the system is larger than the upper bound on efficiency provided by Equation 2.3, i.e.

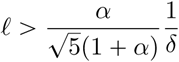

that is, at distance of order 1/*δ*. Consequently, it is impossible to code position with two modules better than this bound.

The question rises then whether the above theoretical bound is achievable, at least for some appropriately chosen *α*. The answer is yes, namely the upper bound in (2.3) is sharp for the golden ratio 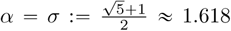

Practically, this also introduces a limit on *∊*(*ℓ*) saying that the phase difference between the modules remains always larger than a specific *lower* bound:

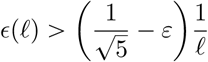

except for a couple of small distances, even for arbitrary small *Ɛ* > 0 (Figure 4b). It may sound strange that there are finitely many exceptions, but in practice these mean only a few instances with *ℓ* being small (Figure 4d). Therefore, if the ratio of the two grid modules equals the golden ratio then the phase difference between the two modules is guaranteed to be larger than the lower bound defined by Equation 2.5. Since *Ɛ* can be arbitrarily small, the lower bound for the golden ratio approaches the theoretical upper bound Equation 2.3 hence σ is an optimal choice for the scale ratio to avoid ambiguity in case of two modules. To give a geometric picture, the golden ratio guarantees approximately uniform coverage of the phase space for both short and arbitrarily large distances (Figure 2c, right).

However, it turns out that there are many good choices: for any algebraic integer *α* of order 2 (i.e. irrational which is a root of a polynomial of degree 2 with integer coefficients) there exists a maximal positive constant *c_α_* > 0 such that

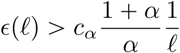

holds except for a couple of small distances (Figure 4c-f, Oxtoby 1980). Hence, the representation is unambiguous whenever *c_α_*(1 + *α*)/(*αℓ*) > (1 + *α*)/(*αδ*), that is up to

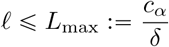

for all *δ* which is small enough.^5^ The constant, *c_α_*, is the single parameter that determines the critical distance up to which encoding is unique and hence can be used to compare different choices of *α* (Figure 4d,f). We already noted that for the golden ratio the lower and the upper bounds coincide (Figure 4b), but the critical distance may be larger for some *α* even if the corresponding lower bound on the phase difference is weaker.

We estimated the value of *c_α_* for various scale ratios at different noise levels. The constant *c_α_* is well defined only for algebraic numbers, but can also be estimated for real numbers from the scaling of the phase difference with distance using numerical simulations (Methods 4.8). Unlike for algebraic numbers, *c_α_* of real numbers depends on the distance range used for the estimation, which we controlled by setting different intervals for *δ* in the simulations (Figure 5).

**FIGURE 5.**
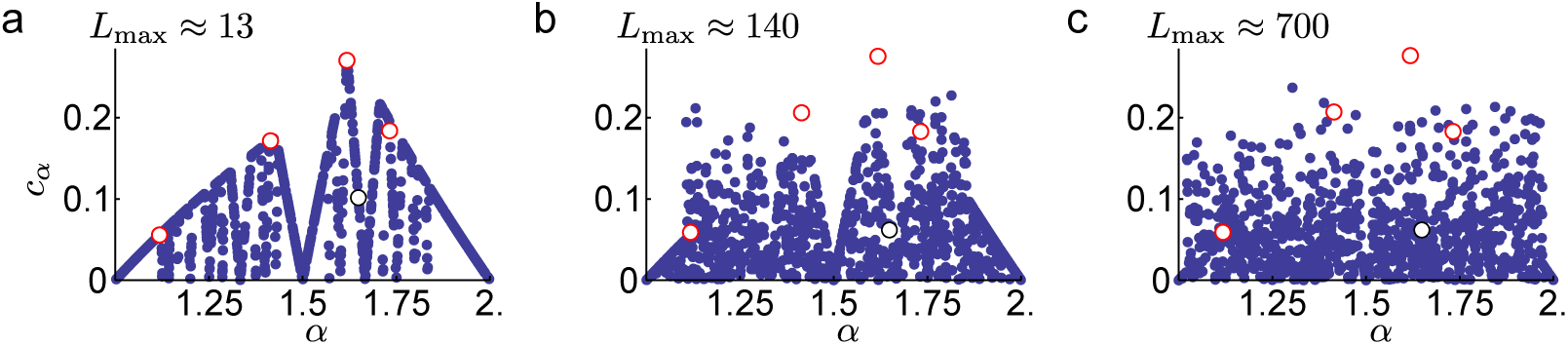
Approximate *c_α_* values as a function of *α*. The values are shown for 1000 *α* randomly selected from the interval [1, 2]. The *c_α_* of algebraic (√2, √3, *σ*, *σ* − 1/2) and non-algebraic (√*e*) irrationals are also shown in red and black, respectively. We estimated c_α_, based on Equation 2.7, as *c_α_* ≈ inf{*L*_max_*δ* | *δ*_min_ < *δ* < *δ*_max_}, for different noise ranges [*δ*_min_, *δ*_max_] in the three panels: (a) 0.005< *δ* < 0.2, (b) 0.005 < *δ* < 0.05, (c) 0.001 < *δ* < 0.01. Approximate *L*_max_ values are indicated on the top of the panels for *α* = *σ*.

These computations confirmed that *σ* is the best scale ratio choice in case of two modules with 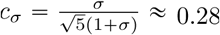 0.28, but also showed that, on both short and long run, *c_α_* is extremely sensitive to the choice of α: in case of a small error in the tuning of *α*, the efficiency can drop substantially and *c_α_* becomes practically 0 (Figure 5), which means that in the immediate neighbourhood of the optimal *α*, there are close to pessimal grid cell configurations. This is because the lower bound on the phase difference (Equation 2.6) requires *α* to be an algebraic number, and in an arbitrary small neighbourhood of any algebraic number there are (infinitely) many non-algebraic numbers, i.e., transcendental numbers (*α = √e*) Figure 4a) or rational numbers (*α* = 3/2, *c_α_* → 0,) Figure 2a). As non-algebraic irrational numbers can be much better approximated with rationals then algebraic numbers, non-algebraic grid scale ratios will lead to much stronger interference between the two modules, but only at distances moderately large compared to the scale of the modules (Figure 5).

The extremely rough landscape of *c*_*α*_ renders optimization for *α* an espacially difficult problem: it is very unlikely that a biological system would be able to find the global optimum for the scale ratio of two grid modules and a relatively small mistuning from a local optimum could significantly deteriorate the efficiency of the system. Therefore, at least in the case of two modules, it seems to be impossible to achieve asymptotically optimal scale ratio for the grid cells. Although the two-module grid cell system was shown to be unstable against small variations in the relative scale of the modules, it is unclear if these results generalise to grid systems with multiple modules.

### 2.3. Multiple modules - naive solution

To derive the general solution for *M* grid modules, we focus on a set of 1-dimensional grids with scales *α*_0_ = 1 < *α*_1_ <… < *α_M-1_*. Spatial representation is unambiguous up to a distance *L* from the origin if there is at least one module for which the phase is significantly different from 0 (Figure 6). Conversely, the representation becomes ambiguous if all modules show interference at the same location, i.e., the phase of all modules are very close to 0 at distance *ℓ* from the origin.

**FIGURE 6.**
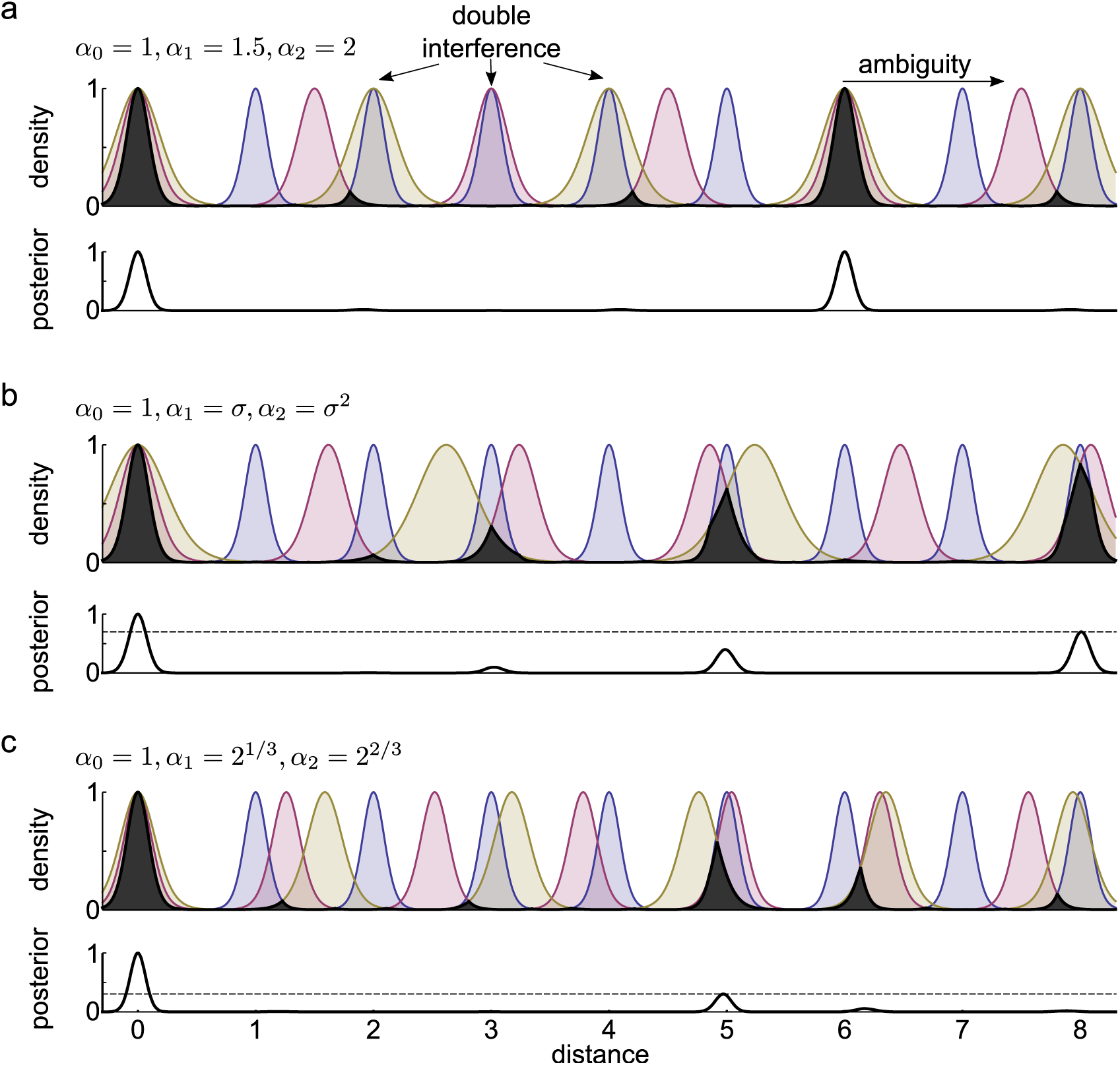
Interference of *M* = 3 modules with different choice of scales. (a) Top: Posterior densities for three modules with rational scale ratios. The overlap between the modules is shown in black, its height indicates the interference of the three modules as a function of distance from the origin. The representation becomes ambiguous only if all 3 modules interfere, as at distance 6. Bottom: Ambiguity in position coding quantified by the multi-modality of the combined posterior. (**b**) Posterior densities for three modules with pairwise optimal scale ratios. The scales are 1 (blue), *σ* (red), and *σ*^2^ (olive). If two modules interfere with each other, then they interfere with the third as well: at distance 8 the three peaks almost coincide. (**c**) The same as in (b) for scales 1, 2^1/3^, 2^2/3^, powers of a cubic algebraic number. Although pairwise interference can be very strong between any pairs (e.g. at distances 5, 6.2 and 8), the total interference is substantially lower than in panel b (bottom).

Theoretical studies demonstrated that geometric progression of grid cell scales is optimal for nested coding, i.e., when each module refines the position coding within the period of the subsequent module (Figure 1c, Mathis et al. 2012a; Wei et al. 2015; Mosheiff et al. 2016). Recordings from multiple grid cells along the dorso-ventral axis of the entorhinal cortex found a set of discrete modules with increasing grid scale (Hafting et al., 2005; Barry et al., 2007; Brun et al., 2008; Stensola et al., 2012). However, the precise geometric progression of grid scales was not confirmed as the scale ratio between the successive modules was found to be highly variable (Stensola et al., 2012). This variability can be partly attributed to the difficulty of estimating the grid scale from finite and noisy data, and the geometric progression could still be a good approximation of the data.

Thus, we speculated whether the set of grid cells with *scale ratio* (*α*) optimally chosen between pairs of successive grid modules were also near optimal in representing spatial location. Such pairwise optimization would lead to a set of scales showing geometric progression with the scale ratio being *α* = *σ*, i.e., [1, *σ*, *σ*^2^,…]. Surprisingly, we found that with scale ratios that are efficient for two modules, adding a third (or more) module provides only very little additional improvement on the coding capacity of the grid cell system.

To see this, consider for example the golden ratio *σ*, which is a second order algebraic number, i.e., it is the root of the integer coefficient polynomial *x*^2^ − *x* − 1. Therefore, the phase *ψ*(*x*) = (*x* mod *σ*^2^)/*σ*^2^ of any spatial point *x* according to the third module can be simply expressed with that of the first two modules as

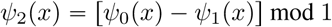

(Methods 4.2). In other words, the phase of the third module provides no additional information given the phase of the other two modules. In particular, if both *ψ*_0_(*x*) and *ψ*_1_(*x*) are close to 0 (Figure 6b), then so is *ψ*_2_(*x*) and hence the third module fails to resolve the ambiguity when the two first modules interfere. The same argument applies for any other low-order algebraic number as well. Nevertheless, in the next section we show that many modules with appropriately chosen scale ratio can perform much better than two modules.

### 2.4. Multiple modules - general solution

The logic of the general solution for multiple modules is the same as in the case of two modules. Here we only state the main results and the technical details of the analysis can be found in the Methods (4.3).

First, we show that a similar upper bound exists for the maximal phase difference between the modules. Compared to the two-module case, the bound is weaker when *M* > 2 as the phase difference scales only with 1/*ℓ*^1/(*M*-1)^ » 1/*ℓ* meaning that it ensures simultaneous interference between all modules only at much larger distances.

Second, we found that the upper bound can be satisfied, up to a constant multiplier, *c_A_* for algebraic scale ratios. Specifically, if the scales of M modules form a geometric series with common ratio *α* being an algebraic number of degree *M*, the upper bound is tight, meaning that the phase difference does not shrink faster than 1/*ℓ*^1/(*M*-1)^. Intuitively, this scaling indicates that there is always at least one pair of modules for which the phase difference at the integer distance *ℓ* from the origin is larger than the lower bound.

The critical distance *L*_max_ up to which coding is unambiguous can be expressed as(cf. Equation 2.7):

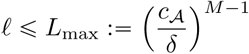

for all *δ* which is small enough. Intuitively, Equation 2.9 demonstrates an exponential scaling of the maximal distance uniquely represented by a population of grid cells with the number of grid modules, *M*. The coding range of a particular set of the grid scales, ***A*** = (*α*_1_,…, *α*_*M* − 1_), depends on both the noise in the system and on the the basis of the exponential *c_A_*

To directly compare the capacity of the non-nested grid cell system derived here with previous estimates, we also calculate *N*_max_, the number of distinguishable spatial phases:

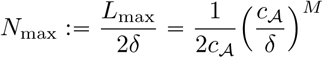

The maximal capacity of the grid cell system has been estimated in the case of nested coding (Wei et al., 2015; Mathis et al., 2012a). Efficient coding with nested modules requires that *α_i_* = *r^i^* with 0 ⩽ *i* ⩽ *M* − 1 and *r* being the scale ratio with fixed relative uncertainty of module 2*δ* = 1/*r* (Wei et al., 2015). The position of the animal can be determined at precision 1/*m* without ambiguity if the animal is restricted to move in an environment with the size identical to the scale of the largest module, *m*^*M*−1^. In this case the number of distinguishable spatial phases is 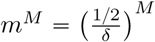 which is identical to the capacity we found for non-nested coding when *c_A_* = 0.5 (Equation 2.10). Since it has been previously shown that the grid system achieves its theoretically maximum capacity for nested codes (Wei et al., 2015; Mathis et al., 2012a), we can conclude that the capacity of the non-nested grid cell system is also nearly maximal, provided that *c_A_* is near 0.5. In the next sections we will first numerically estimate the value of *c_A_* for various choices of the grid scales *A* and then we will show that with sufficiently large number of modules *c_A_* is guaranteed to approach 0.5.

### 2.5 Numerical estimation of the *c_A_*

We developed an efficient method to numerically estimate the value of *c_A_* for various parameter settings that is based on the simultaneous Diophantine approximations of a set of irrational numbers (Methods, 4.8). Using realistic noise levels we found that, in the case of two modules, the efficiency of coding is very sensitive to the choice of *α*, especially to its number theoretic properties (Figure 5a). This sensitivity gradually vanishes with increasing the number of grid modules (Figure 7a-b), and for *M* = 10 modules *c_A_* ∊ [0.2, 0.4] for almost all choices of the grid scales (Figure 7b), both when the scales follow a geometric series with a common scale ratio *α* (Figure 7b) and when all the *M* scales are chosen from the bounded interval [1, 2] (not shown). We also found that *c_A_* vanishes only for pathological examples such as rational numbers or powers of the second order algebraic number *α* = *σ* − 1/2 ≈ 1.118 (Figure 7a-b, red). The only scale choice that significantly degrades the performance is when α ≈ 1 (Figure 7a-b) which means that all grid modules have nearly identical spatial scale.

**FIGURE 7.**
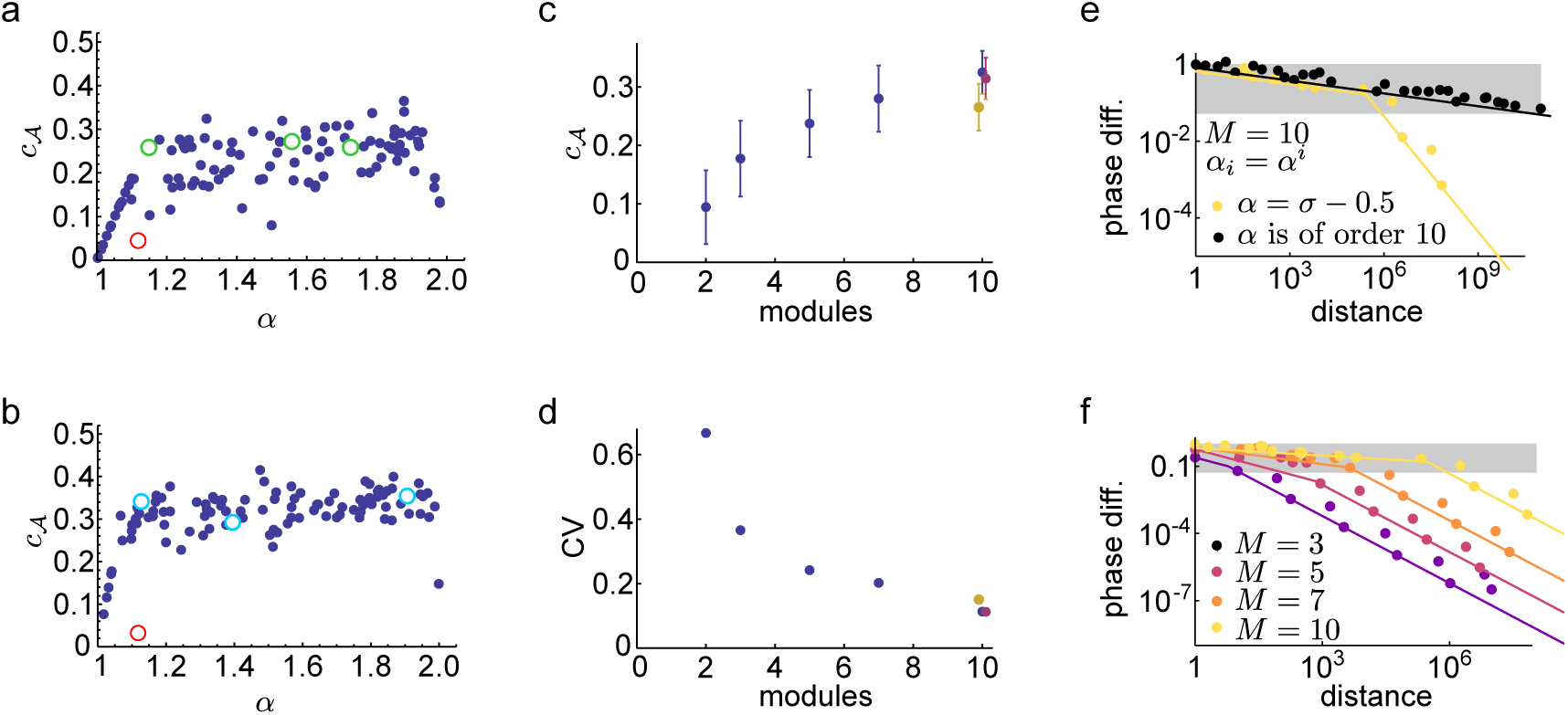
Robustness of the grid code with multiple modules. (a)-(b): Approximate *c*_A_ values estimated for 100 *α* randomly selected from the interval [1, 2] with *M* = 5 (a) and *M* = 10 (b) (see also Figure 5a for *M* = 2). The scales form a geometric series, i.e., 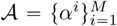 Red circle indicates *c_A_* for a second order algebraic number *α* = *σ* − 1/2 ≈ 1.118 Green (cyan) shows *c_A_* for a 5th (10th) order algebraic numbers. Noise level is the same as in Figure 5a (0.05 < *δ* < 0.2). **(c)-(d)**: Mean (c) and coefficient of variation (d) of *c_A_* evaluated on the range α = {1.1, 1.9} For *M* = 10 the *c_A_* is shown for two alternative selection of the scales: if all 10 scales are selected randomly from the interval [1,2] (olive) and when αs form a geometric series perturbed as 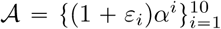 where ε_i_ are i.i.d. uniform random variables on the range [−0.01,0.01] (red). **(e)-(f)** Phase difference (*∊*(*ℓ*), (4.8)) in the function of the distance, *ℓ*. (e) Effect of number theoretic properties with *M* = 10. When *α* is the root of the 10th order polynomial *x*^10^ − *x*^7^ − 1, *α* ≈ 1.12725, *∊*(*ℓ*) decays as 1/*ℓ*^1/9^ (black). When *α* is second order, *α* = σ − 1/2 ≈ 1:1180, the initial decay is similar, but after a critical distance at *ℓ* ≈ 10^6^ the decay becomes 1/*ℓ* (yellow). (**b**) The critical distance grows with the number of modules (*α_i_* = (σ − 1/2)^i^). Grey shading in (e-f) indicates the range of phase difference smaller than noise (Equation 4.9).

To quantify the sensitivity of the grid system to the choice of the scale parameters we calculated the mean and the coefficient of variation of *c_A_* with random choices of (Figure 7c-d). We found, that the average *c_A_* increased monotonically with the number of grid modules indicating that the system’s performance becomes closer to the ideal *c_A_* = 0.5. value as the number of modules increased (Figure 7c). Moreover, the variability of *c_A_* consistently decreased with the number of modules reflecting the improved robustness of the system to the choice of grid periods (Figure 7d). Note, that the maximal distance grows exponentially with the number of modules (Equation 4.10)and *c_A_* ⩽ 0.5 sets the base of the exponential scaling.

To further investigate the mechanisms responsible for the robustness of the system, we numerically evaluated the minimal phase difference between the modules, *∊*(*ℓ*) in the function of the distance (Figure 7e-f). In line with the predictions of the theory (Equation 4.8), we found that the phase difference decreased with *ℓ*^−1/(*M*−1)^ i.e., with a small negative power of the distance for *α* being an order *M* algebraic number (Figure 7e, black). For suboptimal *α*s, the scaling of the phase difference was nearly optimal up to a critical point beyond which the scaling followed the algebraic rank of *α* (i.e., second order *α* scales with 1/*l*, Figure 7e, green). Importantly, this critical point, where the transition occurs between ideal and number theoretical scaling is located at increasingly larger distances when the number of modules increased (Figure 7f). Therefore, the asymptotic, number theoretical properties of the grid periods have a gradually lower impact on the performance of the system in the distance range limited by the intrinsic variability on neuronal spiking (Figure 7ef, background shading).

These observations suggests that even random scale choices might achieve optimal performance as the number of modules grow. In the next section we make this statement mathematically precise and demonstrate that indeed, *c_A_* approaches its maximum, 0.5, when the number of modules grow and the scales are chosen uniformly at random from a bounded interval.

### 2.6. Capacity of non-geometric grid scales

Previous studies claimed that the representational capacity of the grid cell system is exponential in the number of modules (Fiete et al., 2008; Sreenivasan & Fiete, 2011; Mathis et al., 2012a, b; Wei et al., 2015). However, an analytic proof is known only in case of nested coding (Mathis et al., 2012b; Wei et al., 2015), a coding scheme that requires very specific choice for the scaling of the modules. Our number theoretic argument (eq. 4.10) solely does not imply exponential capacity, since it does not exclude the possibility that the base of the exponential *c_A_* converges to 0 as *M* increases (although we observed the opposite trend, see Figure 7c). In this section we investigate the asymptotic properties of the grid code when the number of modules increases and the relative uncertainty *δ* of the modules remains fixed. Here we only state these results informally, and leave the precise statements and the slightly technical mathematical proof to the Methods (4.7).

The main idea behind the proof is that the phase of a given module at particular distance *x* from the origin depends only on the scale of that module, *α*. If the scale is chosen from a bounded interval [1, *α*_max_], then the phase is also random variable with probability distribution approaching the uniform distribution as the distance increases. Then, the probability of simultaneous interference between *M* modules, that is, the probability of all modules being near phase 0 at some distance *x*, is proportional to the volume of an M-dimensional hypercube, which is *V* = (2*δ*)^*M*^, where the side of the cube is 2*δ*. The ratio of the volume of the hypercube and the unit cube (*the number of distinguishable phases*) diminishes exponentially with *M*, and the total distance (expressed in units of *α*^0^ = 1) covered without ambiguity is 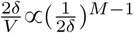 Specifically, our statement is, roughly speaking, that if 0 < *δ* < 1/2 is fixed, *M* is large enough, and the module scales are drawn uniformly at random from a not too narrow bounded interval, e.g. from [1, 2], then the representation is unambiguous up to the exponential distance

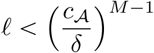

with probability approaching 1, and *c_A_* approaching 1/2. Although this statement is valid only if *M* → ∞ we emphasize that this result is stronger than our previous derivation in three aspects: First, our previous derivation (4.10) allowed *c*_A_ to tend to 0 as *M* increased. Now we showed that this does not happen. In fact, the value of the constant tends to its theoretical maximum 0.5 (section 2.3, Wei et al. 2015) for large *M* with high probability, confirming our previous numerical results (Figure 7c). Second, one can achieve this nearly optimal performance without increasing the scales exponentially, with the scales chosen from a bounded interval. Third, this almost optimal efficiency is not only reached for some appropriately chosen scales, but for almost all choices. Thus, our results demonstrate that no meticulous tuning of the grid scales is required for close to optimal grid system performance.

## 3. Discussion

In this paper we developed a novel analytical technique to investigate the coding properties of grid cells. Using this technique, which is based on Diophantine approximation of real numbers by fractions of integers, we were able to derive several novel and non-trivial properties of the grid cell code. First, we demonstrated that on the long run, the capacity of the system depends heavily and chaotically on the number theoretic properties of the scale ratio between the successive modules. To achieve optimal performance in a system with *M* modules the scale ratio has to be an algebraic number of order *M*. Second, we showed that in the presence of neuronal noise the capacity of the grid code becomes increasingly more robust to the choice of the scale parameters when the number of modules is increased: when *M* > 2, randomly chosen scales perform nearly as well as the optimal scales. Finally, we derived that the capacity of MA and nested grid codes are asymptotically identical (in the large *M* limit), even for randomly chosen scale parameters for the MA codes.

**Information content.** Previous works used specific assumptions to derive exponential information content for the grid cell coding system: they assumed either a nested coding scheme (Mathis et al., 2012b; Wei et al., 2015) or presumed that the phase space is covered evenly and the noise in a given module decreases with the number of modules (Sreenivasan & Fiete, 2011). Here we generalised these findings and demonstrated that nested and MA codes have asymptotically equal capacity.

To derive the capacity of MA codes we realised that achieving uniform coverage of the phase space is not trivial in the case of two or a small number of modules, but can only be attained with appropriately chosen scales. Specifically, we recognised that uniform coverage of the phase space by the phase curve at arbitrary distances is guaranteed if the scale ratio between the two modules is an algebraic number of order 2. Using our formalism allowed us to generalise this intuition for arbitrary number of grid modules and to demonstrate that even a random choice of grid scales guarantees uniform coverage of the phase space when the number of modules is high.

To derive exponential information content we also relaxed the assumption of an earlier studySreenivasan & Fiete (2011) that the total amount of the noise remains constant in the grid system even when the number of modules is increased, i.e., the coding errors of each module decreases with *M*. Here we derived these results using the more realistic assumption that the noisiness of each modules is independent of *M* and proportional to the scale of the module.

We confirmed our analytical results by extensive numerical simulations regarding the simultaneous interference between grid systems with various choices of the scale parameters. In line with previous results (Fiete et al., 2008; Towse et al., 2014), our simulations confirmed that the grid system is robust to the choice of the scale parameter and that the coding range is exponential in the number of modules.

**Nested coding versus MA code.** Although the efficiency of the coding investigated in this paper is slightly worse than that of the optimal nested coding (Mathis et al., 2012b; Wei et al., 2015), MA codes also have several advantages. First it uses orders of magnitude smaller scale lengths than the maximal distance up to which the coding works properly. The largest grid scales measured experimentally are ≈ 3m (Brun et al., 2008), substantially smaller than the typical distances travelled by rodents.

Second, while the consequence of a module failure simple decreases the capacity of the system in the case of MA coding, it can have more dramatic effect in nested codes: Although malfunction of the largest or smallest module reduces either the capacity or the resolution of nested codes, respectively, the lack of intermediate modules functionally breaks the interaction between the remaining modules decreasing both the resolution and the capacity of the system in a disproportionate manner.

Third, once the scales are set, the capacity of nested grid codes does not depend on *δ*, therefore, contrary to MA codes, it is not possible to increase the coding range by inserting more neurons into the same modules. Conversely, the functioning of the nested codes critically depends on accurate decoding of each modules: If the readout neuron does not have access to enough presynaptic neurons from a given module, then the corresponding posterior becomes too wide leading to interference between the modules. This has similar consequences as the absence of the given module in nested codes. In contrast, in MA codes the coding properties remain similar for postsynaptic neurons receiving different number of synapses from different modules, although the coding range is the function of the precision available for the observer. (Equation 4.10).

When encoding dynamic trajectories instead of static location, the number of neurons required to participate in a given module decreases quadratically with the scale of the module, i.e., *n_i_* ∼ 1/*α*_*i*_ (Mosheiff et al., 2016). Specifically, representing the position with a fixed accuracy with *α_i_* = 0.2m requires ∼ 4000 neurons while *α_i_* = 2 m needs only ∼ 40 neurons. This scaling implies that the coding range of the nested grid system can be easily and parsimoniously extended by adding a new module with larger scale but containing only relatively few neurons. Although the relationship between the number of neurons in a module and its scale holds also for MA codes, the total number of neurons required to achieve similar coding range can be substantially smaller in nested codes.

Another consequence of dynamical coding is that the time constant of the readout has to be matched to the scale of the grid modules (Mosheiff et al., 2016). As the grid scale varies over a large range in the case of nested codes, the postsynaptic neuron has to integrate inputs from different grid cells with time constants ranging from 1 ms to 1 second (Mosheiff et al., 2016). In MA codes, the modules have similar scales and their outputs can be integrated with similar time constants.

Finally we note that nested coding and MA coding are not mutually exclusive: they are two extreme forms of decoding the same information, but both can be present in the same system. The MA code has a larger coding range if *c_A_* > *αδ* so it is favoured by small *α* (small differences between scales) and small *δ* (high accuracy). Even in this case locations within the largest grid scale can be decoded as in nested coding, while MA decoder is required beyond this distance.

**Optimization and robustness.** The bewildering regularity of grid cells’ firing fields motivated theories about them being evolved to optimally represent the spatial location of the animal (Kropff & Treves, 2008; Sreenivasan & Fiete, 2011; Mathis et al., 2012a; Moser et al., 2014). Besides the general optimality of triangular grid-like firing fields (Mathis et al., 2015), recent theoretical work derived optimal scale ratio of successive grid modules in the case of nested coding (Wei et al., 2015; Stemmler et al., 2015; Mosheiff et al., 2016). These predictions roughly agree with the average scale ratio observed in the entorhinal cortex (Barry et al., 2007; Stensola et al., 2012; Krupic et al., 2015), but do not explain the substantial amount of variability characteristic of this data.

The optimization principle assumes that substantial improvement in the performance of the system can be achieved with precise tuning of its parameters. In the present study we demonstrated that this is indeed the case in the absence of noise. However, even in this case, optimization would be almost unfeasible for two reasons. First, the coding range is an extremely irregular, discontinuous function of the scale parameter, making optimisation essentially a trial and error game. Second, a scale parameter that is optimal for a given number of modules is guaranteed to be inefficient when the number of modules is increased precluding the possibility of pairwise or modular optimization.

However, taking the variability of neuronal firing into account changes the picture dramatically. We demonstrated that when the coding accuracy of grid modules is limited by neuronal noise, the capacity of the system becomes surprisingly robust to the choice of the scale parameters making its optimization unnecessary. Note, that generating the regular, periodic firing fields of grid cells demands accurate integration of velocity inputs (Issa & Zhang, 2012; Burak & Fiete, 2012) and repeated error correction (Burak & Fiete, 2009; Samu et al., 2009), both requiring the precise tuning of single neuron and network parameters within a given module.

**Predictions.** Our finding, that grid cells have an exponentially large coding range even with randomly chosen grid scales of similar magnitudes makes several important predictions. First, MA coding predicts that the coding range is substantially larger than the largest grid period. Since grid cells are likely to be involved in path integration (McNaughton et al., 2006; Moser et al., 2008), this prediction could be tested by probing path integration abilities of rodents beyond distances of the largest grid period (Etienne & Jeffery, 2004).

Second, in the case of MA coding, different modules have similar contribution to the coding range of the system. Therefore, the effect of targeted dMEC lesion (inactivating a single module, similar to Ormond & McNaughton 2015) on the rat’s navigation behaviour would be largely independent of the actual location of the lesion (i.e., which module is inactivated).

Third, since the performance of the system is independent of the precise choice of the grid scales, we expect a large variability in the scale ratio of successive grid modules both within and across animals. This prediction is consistent with the experimental data available (Barry et al., 2007; Stensola et al., 2012; Krupic et al., 2015), although further statistical analysis would be required to specifically determine the distribution of scale ratios.

Finally, we predict that the performance of the system is not particularly sensitive to changes in the scale parameter of a subset of modules during e.g., global remapping induced by environmental changes (Fyhn et al., 2007). It has been shown that under certain conditions simultaneously recorded grid cells respond coherently within a module and independently across modules to environmental distortions (Stensola et al., 2012). To test the predictions of our theory, the behavioural consequences of incoherent realignment across modules should be assessed and compared with the effects of environmental manipulations inducing coherent realignment (Krupic et al., 2015).

## 4. Methods

### 4.1. Estimating the precision of a single module

We numerically estimated the precision of position coding by a single module by first simulating the motion of the animal as a one dimensional Gaussian random walk:

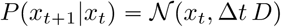

with ∆*t* = 1 ms temporal resolution and *D* = 0.005 m^2^/s, which gives ≈ 5 cm displacement in 0.5 s (Mosheiff et al., 2016). We simulated the activity of *N* = [10,300] grid cells from a single module. Grid cells had a circular tuning curve:

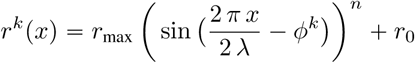

with the following parameters: *r*_max_ = 15Hz, *r*_0_= 0.1Hz, λ = 0.25m and *Φ^k^* chosen to uniformly cover the interval [0, 2π]. The power *n* = 22 was set to match the mean firing rate of the grid cells, 〈*r*(*x*)〉 = 2.5 Hz to experimental data (Fyhn et al., 2007). Larger (λ = 2.5 m) grid spacing was modelled by decreasing the speed of the animal by a factor of 10 (*D* = 0.00005 m^2^/s). The firing rate is shown in Figure 1b, right (olive).

Spike trains were generated as an inhomogeneous Poisson process with neurons conditionally independent given the simulated location:

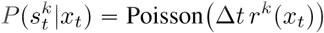

The posterior distribution of the position was numerically calculated by recursive Bayesian filtering:

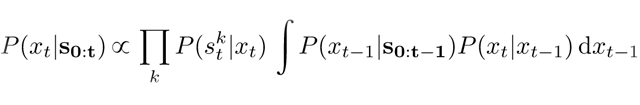

The colormap in Figure 1b shows this posterior distribution with *N* = 50 cells and λ = 0.25 m.

At each timestep the posterior distribution was fitted with a von Mises distribution with a location *μ_t_* and a concentration parameter *κ_t_*. The width of the posterior relative to the grid scale was estimated as:

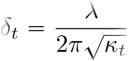

To be conservative, we chose *δ* to be the 99% of the empirical CDF of *δ_t_* The largest *δ* = 0.12 was found with λ = 0.25 m and *N* = 10 cells. The smallest *δ* = 0.01 corresponds to the parameters λ = 2.5 m and *N* = 300 cells.

### 4.2. More than two modules: golden ratio is suboptimal

*Proof of* (2.8). By the definition of the phases *ψ_i_*(*x*) when the animal is at distance *x* from the origin there are some integers *ℓ*, *k*_1_, *k*_2_ so that

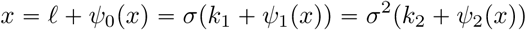

Using that σ^2^ − σ − 1 = 0 we get that

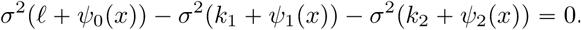

Rearranging terms yields

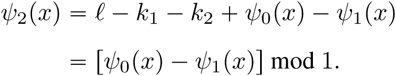

Clearly the same argument works not only for the powers of the golden ratio, but for powers of any algebraic number of order lower than the number of modules.

### 4.3. Interference with *M* modules

To derive the general solution for *M* grid modules, we consider on a set of 1-dimensional grids with scales *α*_0_ = 1 < *α*_1_ <… < *α*_*M* − 1_. Again, the interference between the modules can be expressed by the *simultaneous* Diophantine approximation of the vector 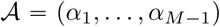 using fractions of integers with the common *numerator ℓ*, i.e., *α_i_* ≈ *ℓ*/*k_i_*. Importantly, a theorem by Dirichlet (Section 4.4) provides an upper bound on the efficiency of the approximation. Namely, for all (*M* − 1)-tuple of irrational numbers *α*_1_,…,*α*_*M* − 1_we have infinitely many collections of integers *k*_0_, *k*_1_,…, *k*_*M* − 1_(with *k*_0_ = *ℓ*), such that the approximation error defined as

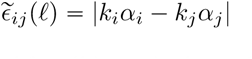

is simultaneously smaller than the upper bound for all items in the tuple:

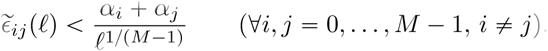

Note, that 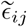 differs from *ε* defined for two modules (Equation 2.2) as it is not normalised with *α*.

For a set of grid scales *α_i_* = *α^*i*^ (*i** = *0,…, *M* − 1)* where *α* is an algebraic number of degree *M*, there exists a maximal positive constant *c_**A**_* such that

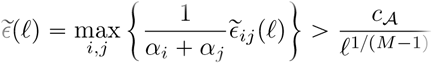

holds, except for at most finitely many integers *ℓ*.

The position representation is unambiguous if there is at least one pair of modules for which the phase difference is larger than the threshold set by the noise, i.e., 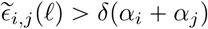 which holds if

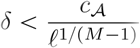

From here, the critical distance *L*_max_ up to which coding is unambiguous can be expressed as (cf. Equation 2.9):

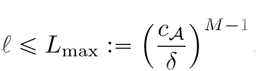

for all *δ* which is small enough. (Section 4.6).

### 4.4 Proof of Dirichlet’s theorem

*Proof of* (4.7). First we prove that any vector of irrationals can be approximated to the claimed order with rationals having the same *denominator*. Let 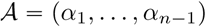 To approximate 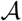 with rationals of denominator at most *Q* let us define the vectors 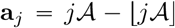, *j* = 0,…, *Q*, where floor is understood coordinate-wise. Let us partition the unit cube [0, 1]^*n*−1^ into small cubes of side length *Q*^−1/(*n*−1)^ so that altogether we have *Q* of them. Since we have *Q* + 1 many a*_j_*-s each falling into [0, 1]^*n* − 1^, hence there will be (at least) 2 of them falling into the same small cube, a*_k_* and a*_l_*, say. Then

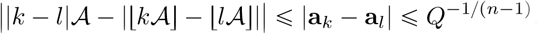

with the inequalities holding coordinate-wise. Therefore, because of | *k* − 1| ≤ *Q*, 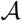 is approximable with denominator |**k* − l*| and numerator (vector) 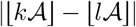 with error not exceeding |*k − l*|^−(1 + 1/(n − 1))^

The desired statement follows then by simultaneously approximating the numbers 1/*α*_i_ with common denominator, which is also a simultaneous approximation of *α_i_* with common numerator, which completes the proof.

### 4.5 Powers of an algebraic number are badly simultaneously approximable

The statement of Drmota & Tichy (1997) (see also Cassels 1955) is that powers of an algebraic number are badly simultaneously approximable with common *denominator* in the following sense. Let *β* be an algebraic number of order *M*. There exists *c*_β_ > 0 such that for all integer *ℓ*, *k_i_* there is *i ϵ* {,…, *M* − 1} for which

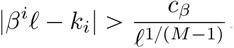

*Derivation of* (4.8). Our goal is to give a lower bound on ❘*α^i^k_i_* − *α^j^k_j_*❘, where *α* is algebraic of order *M*, 0 ≤ *i*, *j* ≤ *M* −1 Without loss of generality suppose that *i* < *j*.

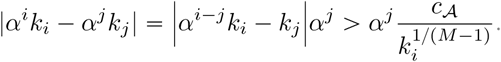

Now the fact that *k_i_* ˜ *ℓ*/α*^i^* implies (4.8) if 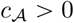 is chosen appropriately.

### 4.6 Coding is unambiguous up to exponential distance in the number of modules

To derive (4.10) we first show that interference of the grid representation is equivalent to pairwise interference between all pairs of modules. To test unambiguity of coding note that the place at distance *x* from the origin is confusable with 0 if for all *i* = 0,…, *M* − 1 there exists an integer *k*_*i*_ such that

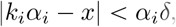

where *δ* is the relative uncertainty of modules. It turns out that, as for *M* = 2, there is no need to consider all *x* ∊ [0, *L*_max_], it is enough to care with integers:

**Claim 4.1**. *There exists x* ∊ [0, *L*_max_] *for which* (4.11) *holds for all i exactly when the following pairwise interference occurs between all modules:*

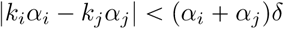

for all *i*, *j* with some integers *k*_i_ (*i* = 0,…, *M* − 1) such that 0 < *k_i_α_i_* ≤ *L*_max_

*Proof*. Let us fix *k*_*i*_, *i* = 0,…, *M* − 1. Pairwise interference means that there is a point *x*_*i*, *j*_ in the intersection of (*k_i_α_i_* − *α_i_δ*, *k_i_α_i_* + *α_i_δ*) = (*a_i_*, *b_i_*) and (*k_j_α_j_* − *α_j_δ*, *k_j_* α + *α_j_δ*) = (*a_j_*, *b_j_*) Due to the topology of the line, it is easy to see by induction that the intersection of all such intervals is nonempty and hence one can chose *x*_*i*,*j*_ = *x*.

The statement is obvious for *M* = 2. Now suppose that the intersection 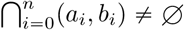 Then it is the interval (*a*, *b*) with

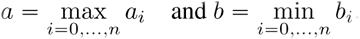

If (*a*_*n*+1_, *b*_*n*+1_) intersects (*a*_*i*_, *b*_*i*_), then both *a*_*n*+1_ < *b_i_* and *b*_*n*+1_ > *a_i_*, and therefore *a*_*n*+1_ < *b* and *b*_*n*+1_ > *a*, which completes the induction. Therefore (4.11) implies (4.12). The other direction is immediate.

Now using Claim 4.1 (4.10) easily follows by rearranging (4.8).

### 4.7. Asymptotic capacity of the random grid cell system

Let us fix the relative uncertainty of modules *δ* < 1/2 and a number *α*_max_ > (1 + *δ*)/(1 − *δ*) We show that if scales *α*_1_, *α*_2_,…: are drawn uniformly at random from [1, *α*_max_], independently of each other, then for any 0 < *ζ* < 1/2 the representation with *M* modules having scales *α*_1_, *α*_2_,…, *α*_*M*_ is unambiguous in every spatial position *x* > 0 up to with probability of order (2*ζ*)^*M*^ as *M* → ∞

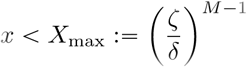

Here *ζ* is the analog of *c*_*A*_ which characterises the capacity of a particular grid cell system. As we will see, the convergence holds for any *ζ* < 1/2 but the speed of the convergence depends on *ζ*: higher efficiency is guaranteed to be achieved only for larger number of modules.

*Proof*: Let *α*_1_, *α*_2_,… be independent random variables distributed uniformly on [1, *A*]. Let *x* be a spatial point and let 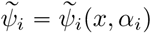 denote the phase of module *i* (with scale *α*_*i*_) at *x*, that is

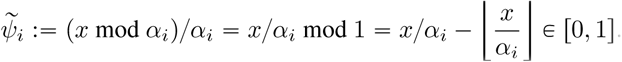

Note that for fixed *x* the distribution of phase 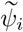 are independent of each other since the α-s are independent. We also use the notation *p*_1_(*x*) for the probability that the phase 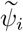 is (almost) indistinguishable from 0, defined in the following way:

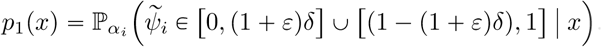

where ε ⇒ 0 is determined later. It is easy to see that *p*_1_(*x*) does not depend on *i*, i.e., it is the same for all modules. Moreover, as the distance increases the distribution of 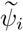 converges to uniform, in particular 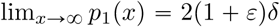 (Figure 8). Hence there exists a critical distance, *x*_0_ = *x*_0_(*δ*, *ε*) for which all *x* > *x*_0_ We have 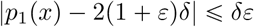 Therefore, for *x* > *x*_0_ we have

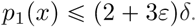

**FIGURE 8.**
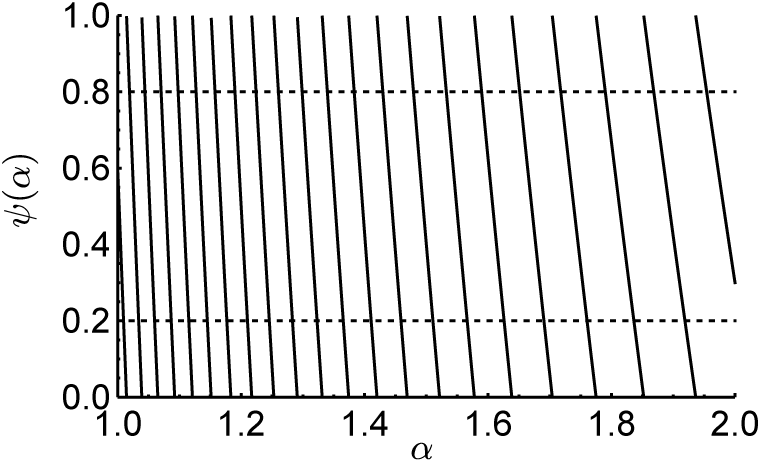
ψ_i_ as a function of *α*_i_ for *α*_max_ = 2, *x* = 42.6 and (1 + ε/3)δ = 0.2. The plot of ψ_i_ consists of line segments more and more vertical as *x* → ∞. Therefore, if *x* is big enough, ψ_i_(*α*_i_) is distributed almost uniformly on [0, 1].

It also implies a bound on the probability of interference of many modules at a given point *x*. If we consider *M* modules with scales drawn uniformly at random from [1, *A*] and independently of each other, then by (4.14) for *x* > *x*_0_ the probability of all phases being close to 0 is where *p_M_*(*x*) is exponentially small in *M*.

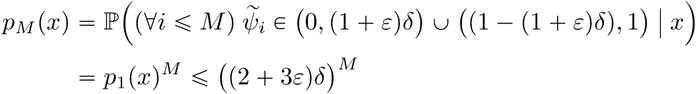

There remains to estimate the probability of interference of many modules anywhere up to a maximally allowed spatial distance. Our goal is to show that

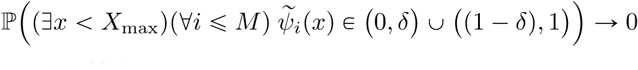

as M → ∞, where 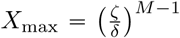 as in (4.13). Note, that satisfying Equation 4.16 is not trivial, since *X*_max_ increases exponentially with *M*.

There is no need to investigate all *x* < *X*_max_ it is enough to show, that there is no interference on a set which is dense enough in [0, *X*_max_] in the stronger sense of (4.15). Indeed, let *Y* be an *ε* dense set in [0, *X*_max_] with at most 2*X*_max_/*ε* elements. Then

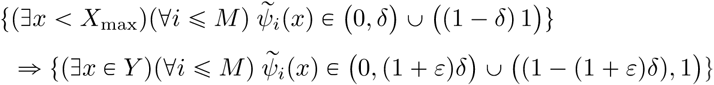

where we used the fact that the 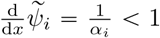 since *α*_i_ was chosen from the interval [1, *A*]. The corresponding inequality for the probabilities of these events is

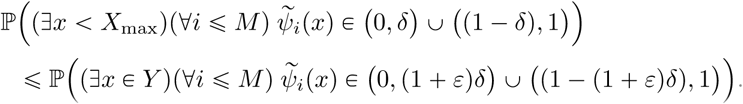

Now for these finitely many points *x* ∊ *Y* we can use (4.15) one by one, if *x*_0_ < *x*:

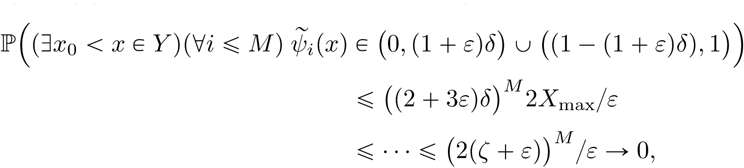

if *ε* < 1/2 - *ζ*, which we assume, where in the first inequality we used (4.15) and union bound, and then in the second one that 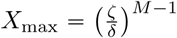 We have to remark that interference in different spatial points is not independent of each other, but union bound works even in that case.

There remains to show that the grid cell representation works up to *x*_0_. Clearly there is no ambiguity up to *x* = 1 + *δ*. To estimate the probability

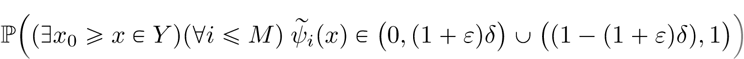

we first have to observe that the cardinality of *Y* ∩ = [1 + *δ*, *x*_0_] is independent of *M*. Therefore to guarantee that the probability in (4.17) goes to 0 we need to show that for all 1 + *δ* ≤ *x* ≤ *x*_0_ there is a scale *α* ∊ [1, *A*] which is able to distinguish *x* from the origin, that is *α* such that

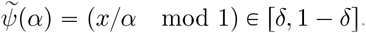

This is so because *x*/*α* is monotonically decreasing in *α* and because

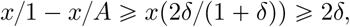

where we used that *α*_max_ > (1 + *δ*)/(1 - *δ*) and *x* > 1 + *δ*. Therefore 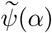 can not lay in [0, *δ*] ∪ [1- *δ*, 1] for all *α* ∊ [1, *α*_max_]

### 4.8. Numerical estimation of the *c_A_* with *M* modules

A common and natural way to numerically investigate Diophantine approximation is using lattice reduction (Lenstra et al., 1982). By lattice we mean a subset 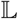 of 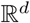 defined by some vectors 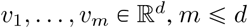 so that

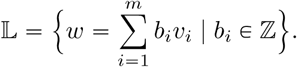

Given a lattice 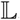 a classical computational problem is to find the shortest non-zero vector of it (Figure 9). In the followings we show how Diophantine approximation of a vector (*α*_1_,…,*α*_*n*_) can be investigated with the help of finding shortest vectors of appropriately chosen lattices.

**FIGURE 9.**
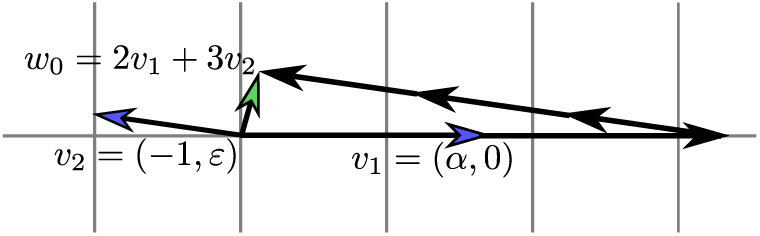
Which element of the lattice generated by the above two blue headed vectors is closest to the origin? Or in other words, what is the shortest nonzero vector which can be obtained as an integer coefficient linear combination of the above vectors?

Let us first consider a simple example. Let the lattice 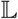 be defined by the rows of the matrix

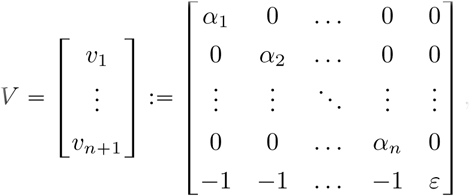

where *ε* > 0. For all *ε* which is small enough the shortest vector 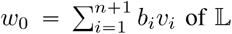 of 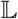 corresponds to a simultaneous Diophantine approximation of (*α*_1_,…,*α*_*n*_) with the common numerator *b*_*n*+1_ and denominators *b*_*i*_, *i* = 1,…,*n*. The parameter *ε* can be considered as a penalty term: the smaller this term the bigger the numerator can be.

When speaking about shortest vectors we need to specify the norm with respect to which vectors are compared. Here we are looking for the largest phase difference between the modules so we use supremum norm (Equation 4.8). The shortest vector in supremum norm of the lattice defined by *V* is an approximation so that

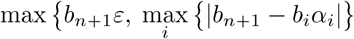

is as small as possible. By this we can compute what is the maximal phase difference between the module with scale 1 and all other modules up to distance *b*_*n*+1_.

Remember that according to (4.8) we are searching for an approximation minimizing

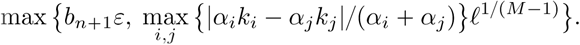

Similarly to the previous example, it can be done simply by dividing columns *i*, *i* = 1,…,*n* of *V* by (1 + *α_i_*) and by adding some more columns of similar form which refer to interference between modules *i* and *j*. For example, for *n* = 3 the shortest (in sup norm) element of the lattice generated by the rows of the following matrix gives an approximation minimizing (4.18):

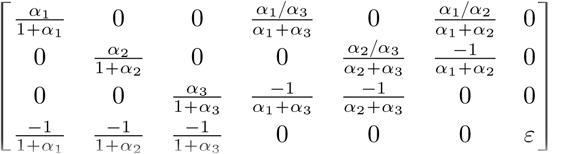

In this way maximal interference in the grid cell system can be computed numerically as shortest vectors of some lattices in supremum norm. Finding this shortest vector is an integer linear programming (ILP) problem, which in general is an NP-hard computational problem, and can be solved by e.g. a branch and bound algorithm (Land & Doig, 1960). There are also efficient methods which find approximation solutions in polynomial time, such as the LLL algorithm due to Lenstra, Lenstra and Lovász (Lenstra et al., 1982).

The LLL algorithm finds not only a short vector of a lattice, but also another basis of it which consists of short and nearly orthogonal vectors in the *L*^2^ norm, a so called LLL reduced basis. The error made by the LLL algorithm is too high to precisely compute the constant terms in (4.10), and therefore we could not rely only on this algorithm. Nevertheless, compared to the ILP solution, we could significantly speed up our computations by first applying the LLL algorithm to find an approximate solution (and a reduced lattice), and then an ILP solver on this LLL reduced basis, which could find nontrivial optimal solutions very efficiently if started from this input.

**Estimation of** 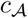. As 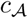 is defined asymptotically (Equation 4.8), in order to estimate it numerically we need an approximation of it for finite distances. An alternative definition of 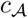 (equivalent with Equation 4.8) is

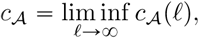

where 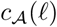 is defined by

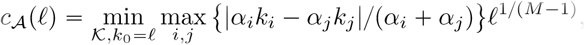

where 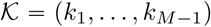 Intuitively, to find the magnitude of interference at location *ℓ*, for all possible values of 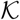 we first select the maximum phase difference in the set and then choose the set with the smallest maximum. The distance versus 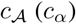 plots of Figure 4 and Figure 7 are indeed distance versus 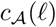 plots. From the plots it is clear that the naive way of approximating 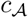 with 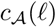 for some large ℓ is not a good idea, as 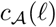 may vary heavily with *ℓ*, especially for non-algebraic scale scale ratios. Note, that estimation of *c*_*α*_ is a special case of with 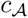 *M* = 2.

To estimate coding efficiency in the presence of noise we are most interested in the above infemum when ℓ is such that the phase difference *c_A_*(*ℓ*)/*ℓ*^1^/(*M*-1) is close to the precision *δ* of the modules. It motivates to investigate the (numerically computable) minimum of the set

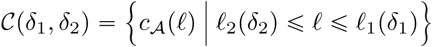

for some pair *δ*_1_ < *δ*_2_, where *ℓ*_2_ is so that for all *ℓ* ⩾ *ℓ*_2_ we have 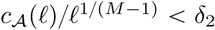 and ℓ_1_ is the smallest ℓ so that 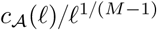. In Figure 5 and Figure 7 the *α* versus 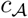 plots are indeed *α* versus 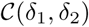 plots, with the *δ*_1_ and *δ*_2_ values given in the captions.

## Acknowledgements

This study was supported by the Hungarian Brain Research Program (KTIA-NAP-12-2-201). We thank G. Orbán and A. Telcs for useful discussions.

### Author Contributions

L.V. and B.B.U. designed the study, L.V. performed the analysis, L.V. and B.B.U. discussed the results and B.B.U. wrote the paper with help from L.V.

## Footnotes

3 Note that *∊*(*ℓ*) can not be the phase difference *in general*, it is the phase difference *at integer distance ℓ*. The phase difference being zero does not necessarily imply interference, only if the phase is also close to 0. In Fig 2, both grids are around phase 0.4 at the distance 2.4 without ambiguity.

4 Note that this argument is correct only if the phase representation ambiguity of modules is independent of the actual position, which holds if we suppose that firing fields of cells from the same module are spaced evenly, which we do assume. Therefore, strong grid codes require uniform coverage of the phase space across arbitrary distances (Sreenivasan & Fiete, 2011).

5 This last condition is only needed to exclude the possible exceptional ℓ distances in (2.5), which in practice is not a crucial condition (Figure 4d-e).

